# Acute Hepatitis E Virus infection in two geographical regions of Nigeria

**DOI:** 10.1101/178756

**Authors:** I.M. Ifeorah, T. O. C. Faleye, A. S. Bakarey, M. O. Adewumi, A. Akere, E. C. Omoruyi, A. O. Ogunwale, J. A. Adeniji

## Abstract

Hepatitis E virus (HEV) remains a major public health concern in resource limited regions of the world. Yet data reporting is suboptimal and surveillance system inadequate. In Nigeria, there is dearth of information on prevalence of acute HEV infection. This study was therefore designed to describe acute HEV infection among antenatal clinic attendees and asymptomatic community dwellers from two geographical regions in Nigeria.

In this study 750 plasma samples were tested for HEV IgM by Enzyme Linked lmmunosorbent Assay (ELISA) technique. The tested samples were randomly selected from a pool of 1,115 samples previously collected from selected populations (pregnant women – 272, Oyo community dwellers – 438, Anambra community dwellers – 405) for viral hepatitis studies between September 2012 and August 2013.

One (0.4%) pregnant woman in her 3^rd^ trimester had detectable HEV IgM, while community dwellers from the two study locations had zero prevalence rates of HEV IgM.

Detection of HEV IgM in a pregnant woman, especially in her 3^rd^ trimester is of clinical and epidemiological significance. The need therefore exists for establishment of a robust HEV surveillance system in Nigeria, and especially amidst the pregnant population in a bid to improve maternal and child health.

## Introduction

Hepatitis E Virus (HEV), is a small, non-enveloped spherical particle of about 32-34nm in diameter and has a single stranded, positive sense RNA genome of approximately 7.5kb that is surrounded by an icosahedral capsid ^1^. It is the only member of the genus Hepevirus and has four genotypes capable of causing human infection.^2,3^Hepatitis E virus (HEV) infection is a major public health concern especially in sub-Saharan Africa among other developing countries.^4,5^ It is one of the most common causes of acute hepatitis and jaundice in the world and has affected about one-third of the world’s population.^6^ Annually, about 20 million new cases and 33 million acute cases of HEV infection occur globally and HEV-related hepatic failure is responsible for approximately 56,600 deaths per year.^7,8^

While it is established that HEV transmission occurs predominantly via the fecal-oral route^9,10^, parenteral, person-to-person, perinatal mode of transmission and eating undercooked wild boar, deer and pork have also been suggested.^11-14^ In developing countries, infection occurs both sporadically and as an epidemic, affecting a large proportion of the population.^15^

In human population, prevalence of anti-HEV varies from one population to another, with significant increase noted among pregnant women and children <2yearrs old.^4^ HEV infection is usually asymptomatic and self-limiting^16^, however, in some cases, acute infection may develop to fulminant hepatitis with high mortality especially among pregnant women in their third trimester^17^,even as chronic forms of HEV have been documented among different populations.^18-^ 20 Diagnosis of HEV infection is based on detection of anti-HEV IgM, anti-HEV lgG and HEV RNA. Specifically, the presence of anti-HEV IgM is a marker of acute HEV infection.^21^

Despite, the significance of HEV in public health, especially in resource limited regions of the world, data reporting remained suboptimal, consequent of inadequate surveillance system.^22^ In Nigeria, there exists dearth of information on prevalence and circulation of HEV infectionin the population. In a bid to bridge this knowledge gap, we attempted, in this study, to describe acute HEV infection among selected population groups in two geographical regions in Nigeria.

## Methodology

Seven hundred and fifty (750) plasma samples were randomly selected from a pool of 1,115 samples previously collected for viral hepatitis studies. The 1,115 plasma samples were collected from consenting apparently healthy community dwellers and ante-natal clinic attendees who participated in our previous studies conducted between September 2012 and August 2013. Specifically, the study populations include: (1) pregnant women (n=272) attending antenatal clinic in two different hospitals in Ibadan, Oyo state, Southwest, Nigeria, (2) asymptomatic community dwellers (n=438) residing in and around Ibadan, Oyo state, Southwest, Nigeria and (3) asymptomatic community dwellers (n=405) from three selected communities in Anambrastate, Southeast Nigeria (Figure 1). These samples were stored at -86 °C at the Department of Virology, College of Medicine, University of Ibadan, Nigeria. A total of 250 plasma samples per population were randomly selected to make the 750 samples analyzed in this study.

**Figure 1:**
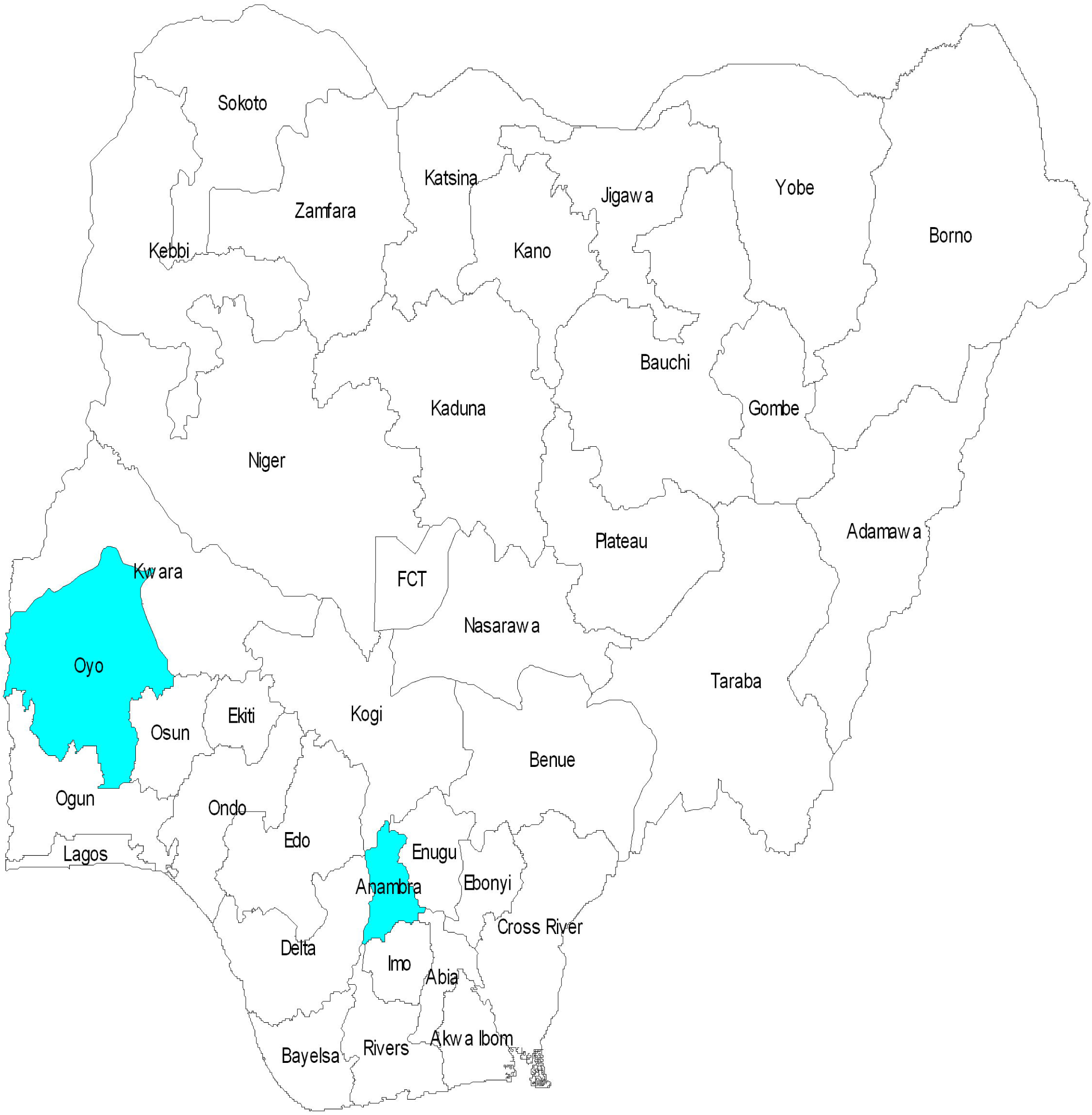
Map of Nigeria showing the Study Communities.

All samples were screened for HEV IgM using Enzyme Linked lmmunosorbent Assay (ELISA) kits (Wantai Biological immunoassay Beijing China) with documented sensitivity and specificity of 97.10% and 98.40%, respectively. Assays were performed according to manufacturer’s instructions. Optical density was read using the Emax endpoint ELISA microplate reader (Molecular Devices, California, USA) and the results interpreted accordingly. Demographic and other relevant information were obtained from the participants using a well-structured questionnaire. Ethical approval for the study was granted by Ul/UCH Ethics committee (UI/UCH/1 1/0058), Oyo State Ministry of Health (A03/479/349) and Anambra State Ministry of Health (MH/PHD/MISC/1)

## Results

In the pregnant women cohort (age range: 17-43 years;median age: 29 years), one (0.4%) participant had detectable HEV IgM. Majority (50%) of the pregnant women, including the only positive participant are within age group 31-40 years. Further analysis shows 2%, 44.8% and 3.2% for age groups 15-20, 21-30 and 41-45 years, respectively. Forty-four (17.6%), 74 (29.6%) and 132 (52.8%) of the pregnant women were in their 1^st^, 2^nd^ and 3^rd^ trimester of gestation, respectively. It is noteworthy that the participant with detectable HEV IgM was in her 33^rd^ week (3^rd^ trimester) of gestation (Table 2).

**Table 1:** Characteristics and HEV IgM prevalence among study participants.

| Variable | Ibadan |  | Ibadan |  | Anambra |  |
| --- | --- | --- | --- | --- | --- | --- |
|  | pregnant women |  | general population |  | general population |  |
| Age | Frequency | Percentage (%) | Frequency | Percentage (%) | Frequency | Percentage (%) |
| 20 and less | 5 | 2.00 | 28 | 11.2 | 41 | 16.4 |
| 21 – 30 | 112 | 44.8 | 81 | 32.4 | 64 | 25.6 |
| 31 – 40 | 125 | 50.0 | 74 | 29.6 | 51 | 20.4 |
| 41 – 50 | 8 | 3.20 | 37 | 14.8 | 47 | 18.8 |
| 51 and above | --- | -- | 30 | 12.0 | 47 | 18.8 |
| Sex |  |  |  |  |  |  |
| Male | --- | -- | 73 | 29.2 | 113 | 45.2 |
| Female | 250 | 100 | 177 | 70.8 | 137 | 46.8 |
| Total | 250 | 100 | 250 | 100 | 250 | 100 |
| HEV IgM |  |  |  |  |  |  |
| Positive | 1 | 0.40 | 0 | 0 | 0 | 0 |
| Negative | 249 | 99.6 | 250 | 100 | 250 | 100 |
| Gestation age |  |  |  |  |  |  |
| 1<sup>st</sup> trimester | 44 | 17.6 | -- | -- | -- | -- |
| 2<sup>nd</sup> trimester | 132 | 52.8 | -- | -- | -- | -- |
| 3<sup>rd</sup> trimester | 74 | 29.6 | -- | -- | -- | -- |

**Table 2:** HEV acute infection among pregnant women.

| Age | 1<sup>st</sup> trimester |  |  | 2<sup>nd</sup> trimester |  |  | 3<sup>rd</sup> trimester |  |  |
| --- | --- | --- | --- | --- | --- | --- | --- | --- | --- |
|  | No | No | No | No | No | No | No | No | No |
|  | tested | positive | negative | tested | positive | negative | tested | positive | negative |
|  |  | (%) | (%) |  | (%) | (%) |  | (%) | (%) |
| 20 and less | 1 | Nil | 1 (100) | 3 | Nil | 3 | 1 | Nil | 1 |
| 21 – 30 | 18 | Nil | 18 (100) | 53 | Nil | 53 | 41 | Nil | 41 |
| 31 – 40 | 22 | Nil | 22 (100) | 71 | Nil | 71 | 31 | 1 | 30 |
| 41 – 50 | 3 | Nil | 3 (100) | 5 | Nil | 5 | Nil | Nil | Nil |

Overall, the Ibadan community dwellers (age range: 1.5-87 years; median age: 40 years), and the Anambra residents (age range: 15-70years; median age: 28.4 years) had zero prevalence rate for HEV IgM. Altogether, 70.8% and 29.2% of the Ibadan community dwellers were females and males, respectively. Further, highest (25.6%) and lowest (16.6%) frequency were recorded among age groups 21-30 and ≤ 20 years, respectively. For the Anambra residents, 54.8% were females, while 45.2% were males. Also, highest (29.6%) frequency was recorded in age group 21-30 years, while, age groups ≤20 had the lowest (0.05%) (Table 1).

## Discussion

In this study, we report HEV IgM rate of 0.4% in the pregnant women cohort, and zero prevalence rate among asymptomatic community dwellers in two different geographical regions of Nigeria. Our finding is in congruence with the rates of 0.9% reported in a study among different populations including apparently healthy individuals in Plateau state^23^, but differs from 1.7% reported among ARV naïve and experienced HIV patients in Ibadan.^24^ Though, varied rates of HEV IgM have been reported in other endemic regions of the world among different populations.^25-27^Thereasons for this discrepancy may range from variation in populations studied and test kits used among others.

Furthermore wide variations in reported seroprevalence of HEV antibodies in Africa during non-epidemic periods of acute and symptomatic hepatitis have been documented.^28^Additionally, studies have indicated that commercially available ELISA kits for anti-HEV antibodies differ dramatically in their sensitivity and specificity.^29-31^ In our study we used an ELISA kit that has been confirmed to have the highest diagnostic sensitivity and specificity when compared to many others by several studies.^30,32^ Specifically, Vollerman *et al.,* (2016) in a comparative study to monitor HEV antibody seroconversion in asymptomatically infected blood donors using 9 different commercially available ELISA kits reported the best performance with the kit used in this study.^33^ Particularly, considerable variation for detection period of IgM antibodies was observed. Therefore, this implies that the prevalence rates of anti-HEV IgM recorded in this study are most likely true representative for the study population as at the time of testing.

Results of this study might imply that the studied communities have good supply of clean water, as drinking of fecal contaminated water has been implicated as one of the major route of HEV transmission.^34^ Further, considering that HEV is known to cause acute infection^4^, a study among asymptomatic persons may not capture acutely infected individuals who might be too sick to participate in such studies. However, this report does not underestimate the burden of acute HEV infection, especially, in resource limited countries as recent reports have confirmed outbreaks with dire consequences in different parts of sub-Saharan Africa including the Northern part of Nigeria.^35,36^This therefore calls for the establishment of a robust surveillance system with focus on HEV that currently seems to be neglected.

The rate of 0.4% found among the pregnant women is in agreement with the rates of 0.5%,0.6% and 0.67% reported among pregnant women in different regions of the world including Africa.^37-39^ Our finding, though higher than 0% rate reported in a similar cohort in France,^40^ is lower than rates ranging from 2.6%-33% recorded by several authors in a similar cohort in different regions of the world.^41-45^ Variation in the performance of different ELISA kits employed in these studies may account for the difference in rates reported.

The fact that the only pregnant woman with HEV IgM in this study was in her 33^rd^ week of gestation, signifies a recent exposure to HEV and the medical significance of the case. The reason being that acute HEV in pregnancy can progress to fulminant hepatitis with a high mortality rate, especially, if it occurs in the 3^rd^trimester.^17^Further, reports have shown that HEV infection during pregnancy can lead to maternal mortality rate of 15% to 25%, especially, with genotype 1 which together with genotype 2 are prevalent in the developing countries.^46^

Acute HEV infection during pregnancy is associated with dire consequences for the unborn, including high rate of miscarriage, stillbirth, neonatal death, high risk of vertical transmission and high risk of preterm delivery with poor neonatal survival rates.^47,48^ It was reported that HEV infection might be responsible for 2,400 to 3,000 stillbirths each year in developing countries, with many additional fetal deaths linked to antenatal maternal deaths.^49^ It is worth mentioning that we do not have information on the outcome of this singular woman’s pregnancy. Hence, we could not document the impact of the HEV infection in the 3^rd^ trimester of pregnancy on either the mother or child.

Recent outbreaks of HEV infection have been documented among Nigerian refugees in Chad and in the Northern part of Nigeria with a significant number of pregnant women being affected.^35,36,50^ With all the health challenges posed by acute HEV infection (especially among pregnant women) it is still grossly under reported in developing countries including Nigeria. For example, there are no laid down policies on HEV management in the country. The pregnant woman with detectable HEV IgM has documented report of fever and jaundice within the preceding 2 weeks prior sampling. Thus, it is probable that the symptoms were as a result of HEV infection, since the woman tested negative to other viral infections with the possibility of showing similar symptoms^51^. Laboratory screening for acute HEV infection is rarely done in Nigeria and more scarcely among pregnant women even when presenting with symptoms suggestive of hepatitis infection (especially when there is no outbreak). Routinely, HBV is the only hepatitis virus pregnant women are screened for in Nigeria. Therefore, it is safe to assume that screening for HEV IgM was not carried out for this woman at the ante-natal clinic. Thus, she obviously might not have been managed for HEV infection. This brings to light how acute HEV infections are possibly missed despite its clinical and epidemiological implications for the subject and the population, respectively.

## Conclusion

We documented acute HEV infection in a pregnant woman in Ibadan, Oyo State, Southwest Nigeria, and reported zero prevalence rates of acute HEV infection among asymptomatic community dwellers in two different locations in the country. Consequently, we recommend more extensive studies on HEV infection in the population in a bid to define the dynamics of this neglected viral infection. In addition, we recommend the introduction of HEV IgM screening for pregnant women in Nigeria. The public should be educated on HEV, its modes of transmission and intervention strategies including vaccination in accordance with the WHO recommendations for safe vaccine.^52^

## Acknowledgement

The authors thank the participants enrolled in this study and extend our sincere appreciation to Professor O. G. Ademowo of the Institute for Advanced Medical Research and Training (IAMRAT), College of Medicine, University of Ibadan, for providing Laboratory space and equipment used at some point during the execution of this study.

This study was carried out by the Evolutionary Dynamics of Hepatitis in Nigeria (EDHIN) research group. This study was not funded by any organization.

### Authors’ contributions

a. Study Design (IIM, FTOC, AMO, BSA)
b. Sample collection (IIM, AMO, OEC,AA)
c. Reagent acquisition, Laboratory and Data analysis (All authors)
d. Wrote first draft of the manuscript (IIM and FTOC)
e. Revised the manuscript (All authors)
f. Read and approved the final draft (All authors)

## References

1. Emerson SU, Purcell RH: Hepatitis E Virus. In Fields Virology. Fifth Edition. Edited by Knipe DM, Howley PM. Philadelphia: Lippincott Williams and Wilkins; 2007:3047–3058.

2. Lu L, Li C, Hadedorn CH. Phylogenetic analysis of global hepatitis E Virus sequences: genetic diversity, subtypes and genetic diversity, subtypes and zoonosis. Rev Med Virol. 2006;16(1):5–36.

3. Sridhar S, Lau SK, Woo PC. Hepatitis E: A disease of re-emerging importance. J Formos Med Assoc 2015; 114: 681–690.

4. Teshale EH, Hu DJ: Hepatitis E: epidemiology and prevention. World J Hepatol. 2011; 3: 285–291

5. Kamar N, Bendall R, Legrand-Abravanel F, Xia NS, Ijaz S, Izopet J, Dalton HR. Hepatitis E. Lancet. 2012;379: 2477–2488.

6. Kim Jfl, Nelson KE, Panzner U, Kasture Y, Labrique AB, Wierzba TF. A systematic review of the epidemiology of hepatitis E virus in Africa. BMC Infect Dis. 2014; 14: 308

7. WHO. Hepatitis E, WHO fact sheet: No. 280, updated July 2015. Available from: URL: http://www.who.Int/mediacentre/factsheets/fs280/en/

8. Bazerbachi F, Haffar S, Garg SK, Lake J R. Extra-hepatic manifestations associated with hepatitis E virus infection: a comprehensive review of the literature. Gastroenterol Rep (Oxf) 2016; 4: 1–15.

9. Guthmann JP, Klovstad H, Boccia D, Hamid N, Pinoges L, Nizou JY, Tatay M, Diaz F, Moren A, Grais RF, Ciglenecki I, Nicand E, Guerin PJ. A large outbreak of hepatitis E among a displaced population in Darfur, Sudan, 2004: the role of water treatment methods. Clin Infect Dis 2006, 42:1685–1691.

10. Howard CM, Handzel T, Hill VR, Grytdal SP, lanton C, Kamili S, Drobeniuc J, Hu D, Teshale E: Novel risk factors associated with hepatitis E virus infection in a large outbreak in northern Uganda: results from a case control study and environmental analysis. Am J Trop Med Hyg. 2010; 83:1170–117.

11. Boxall E, Herborn A, Kochethu G, Pratt G, Adams O, ljaz S, Teo C-G: Transfusion-transmitted hepatitis E in a “nonhyperendemic” country. Transfus Med. 2006; 16:79–83.

12. Matsubayashi K, Kang J-H, Sakata H, Kazuaki Takahashi K, Shindo M, Kato M,Sato S, Kato T, Nishimori H, Tsuji K, Maguchi H, Yoshida J-i, Maekubo H, Mishiro S, Ikeda H: A case of transfusion-transmitted hepatitis E caused by blood from a donor infected with hepatitis E virus via zoonotic food-borne route. Transfusion. 2008; 48:1368–1375.

13. Teshale EH, Grytdal SP, Howard C, Barry V, Kamili S, Drobeniuc J, Hill VR, Okware S, Hu D J, Holmberg SD: Evidence of person-to-person transmission of hepatitis E virus during a large outbreak in Northern Uganda. Clin Infect Dis. 2010; 50:1006–101.

14. El-SayedZaki M, Abd El Aal A, Badawy A, El-Deeb DR, El-Kheir NYA: Clinico-laboratory study of mother-to-neonate transmission of hepatitis E virus in Egypt. Am J Clin Path 2013; 140:721–726.

15. RakeshAggarwal. Hepatits E: Epidemiology and Natural History. J Clin Exp Hepatol. 2013 Jun; 3(2):125–133.

16. Fix AD, Abdel-Hamid M, Purcell RH, Shehata MH, Abdel-Aziz F, Mikhail N, el Sebai H, Nafeh M Habib M, Arthur RR, Emerson SU, Strickland CT. Prevalence of antibodies to hepatitis E in two rural Egyptian communities. Am J Trop Med Hyg, 2000; 62:519–523.

17. Stoszek SK, Abdel-Hamid M, Saleh DA, El Kafrawy S, Narooz S, Hawash Y, et al. High prevalence of hepatitis Eantibodies in pregnant Egyptian women. Trans R Soc Trop Med Hyg. 2006; 100:95–101.

18. Gonzalez Tallion AI, Moreira Vincente V, MateosLindemann ML, Achecar Justo LM. Chronic hepatitis E in an immunocompetent patient. GastroenterolHepatol. 2011;34(6):398–400.

19. Scholosser B, Stein A, Neuhaus R et al. Liver transplant from a donor with occult HEV infection induced chronic hepatitis and cirrhosis in the recipient. J. Hepatol. 2012; 56 (2):500–502.

20. Koenecke C, Pischke S, Hein A et al. Chronic hepatitis E in hematopoietic stem cell transplant patients in a low endemic country? Transpl Infect Dis. 2012;14(1):103–106.

21. NassimKamar, Harry R. Dalton, Florence Abravanel and Jacques Izopet. Hepatits E virus infection Clin Microbiol Rev. 2014 Jan;27(1):116–138.

22. Rayis DA, Jumaa AM, Gasim GI, Karsany MS, Adam I: An outbreak of hepatitis E and high maternal mortality at Port Sudan, Eastern Sudan. Pathog Glob Health. 2013; 107:66–68.

23. Surajudeen A. Junaid, Samuel E. Agina and Khadijah A. Abubakar. Epidemiology And Associated Risk Factors Of Hepatitis E Virus Infection In Plateau State, Nigeria.Virology: Research And Treatment 2014:5 15–26.

24. Georgina N. Odaibo and David O. Olaleye. Hepatitis E virus infection in HIV positive ART Naïve and Experienced Individuals in Nigeria. World Journal of AlDS, 2013; 3: 216–220.

25. El Sayed ZM, Othman W: Role of hepatitis E infection in acute on chronic liver failure in Egyptian patients. Liver Int 2011, 31:1001–1005

26. Farshadpour F, Taherkhani R, Makvandi M. Prevalence of Hepatitis E Virus among Adults in South-West of Iran. Hepat Res Treat 2015;

27. Kuan A. Traoré, Jean BienvenueOuoba, Hortense Rouamba, Yacouba K. Nébi é, Honorine Dahourou, Frédéric Rossetto, Alfred S. Traoré, Nicolas Barro, Pierre Roque. Hepatitis E Virus Prevalence among Blood Donors, Ouagadougou, Burkina Faso. Emerging Infectious Diseases. April 2016;22:(4) 775-776. • www.cdc.gov/eid •

28. Massimo De Paschale, Cristina Ceriani, Luisa Roman,TeresaCerulli, Debora Cagnin, Serena Cavallari, Joseph Ndayake, DieudonneZaongo, KoumaDiombo, Gianbattista Priuli, Paolo Vigan and Pierangelo Clerici. Epidemiology of hepatitis E Virus Infection During Pregnancy In Benin. Tropical Medicine and International Health. jan 2016;21(1): 108–113.

29. Drobeniuc J, Meng J, Reuter G, Greene Monfort T, Khaudyakova N, Dimitrova Z, Kamili S, Teo CG. Serologic assays specific to immunoglobulin M antibodies against hepatitis E virus: pangenotypic evaluation of performances. Clin Infect Dis 2010: 51: e24–e27.

30. Jurgen Wenzel J., Julia Preiss, Mathias Schemmerer, Babara Huber, Wolfgng Jilg. HEV IgG Assays Strongly Influence Hepatitis E Seroprevalence Estimates. J Infect Dis. 2013; 207 (3): 497–500.

31. Ashish C Shrestha, Robert L.P. Flower, Clive R Seed, Susan L Stramer, and Helen M Faddy A Comparative Study Of Assay Performance Of Commercial Hepatitis E Virus Enzyme Linked Immunosorbent Assay Kits In Australian Blood Donor Samples. J. Blood Transfus. 2016; 9647675

32. Pas SD, Streefkerk RH, Pronk M, De Man RA, Beersma MF, Ostertaus A D, Van der Ejik A. Diagnostic Performance Of Selected Commercial HEV IgM And IgG Elisa For Immunocompromised And Immunocompetent Patients. J Clin Virol. 2013 Dec;52(4) 629–34.

33. TanjaVollerman, Juergen Diekman, Matthias Eberhardt, CornrliusKnabbe and Jens Dreier. Monitoring Of Anti-Hepatitis E Virus Antibody Seroconversion In Asymptomatically Infected Blood Donors: Systematic Comparison Of Nine Commercial Anti-HEV IgM And IgG Assays. Viruses. 2016 Aug;8(8):232

34. Aggarwal R, Kini D, Sofat S, Naik SR, Krawczynski K. Duration of viraemia and faecal viral excretion in acute hepatitis E. Lancet 2000; 356:1081–1082.

35. ProMED-mail27jun2017. http://outbreaknewstoday.com/nigeria-e-hepatitis-e-borno-state-cholera-kwara-state-84944/

36. ProMED-mail 26th May 2017. https://www.voaafrique.com/a/au-moins-30-morts-de-lhepatite-e-dans-e-sud-est-du-niger/3872514.html

37. Lindemann MLM, Gabilondo G, Romero B, de la Maza OMS, Perez-Gracia MT. Low prevalence of hepatitis E infection among pregnant women in Madrid, Spain. Journal of medical virology. 2010; 82(10):1666–8

38. GuangyuGu, Hongyu Haung, LeZang, YongchunBi, Yali Hu, Yi-Hua Zhou. Hepatitis E virus Seroprevalence in Jiangsu, China, and postpartum evolution during six years. BMC infect.Dis. 2015;15:560.

39. Abebe M, Ali I, Ayele S, Overbo J, Aseffa A, Mihret A (2017) Seroprevalence and risk factors of Hepatitis E Virus infection among pregnant women in Addis Ababa, Ethiopia. PLoS ONE 12(6):e0180078

40. Renou C, Gobert V, Locher C, Moumen A, Timbely O, Savary J, Roque-Afonso A. Prospective study of Hepatitis E Virus infection among pregnant women in France. Virol J. 2014; 11:68.

41. Adjei AA, Tettey Y, Aviyase JT, Adu-Gyamfi C, Obed S, Mingle JA, et al. Hepatitis E virus infection is highly prevalent among pregnant women in Accra, Ghana. Virol J. 2009; 6:108.

42. Charles M Goumba, Emmanuel R, Yandoko- Nakouné and Narcisse P Komas. A fatal case of acute hepatitis E among pregnant women, Central African Republic. BMC Research Notes 2010, 3:103.

43. Haung F, Ma T, Li L,Zeng W, Jing S. Low seroprevalence of hepatitis E virus in Pregnant women in Yunnan, China. Braz J infect.Ds. 2013;17:716–7.

44. Ahumibe AA, Okonkwo G, Shehu S.M., Waziri E.N, Nguk P. Prevalence of hepatitis E markers in primary health care antenatal attendees in Abuja, Nigeria.16th ICID Abstracts / International Journal of Infectious Diseases 21S (2014) 397.

45. Cong W, Sui JC, Zhang XY, Qian AD, Chen J, Zhu XQ. Seroprevalence of hepatitis E virus among Pregnant women and control subject in China. J. Med. Virol. 2015; 87: 446–50.

46. Ranger-Rogez S, Alain S, Denis F. Hepatitis viruses: mother to child transmission [article in French]. Pathol Biol (Paris) 2002;50(9):568–75.

47. Ornoy A, Tenenbaum A. Pregnancyoutcome following infections by coxsackie, echo, measles, mumps, hepatitis, polio and encephalitis viruses. Reprod Toxicol. 2006;21(4):446–57.

48. Krain LJ, Atwell JE, Nelson KE, Labrique AB. Fetal and neonatal health consequences of vertically transmitted hepatitis E virus infection. Am J TropMed Hyg.2014;90(2):365–70.

49. Patra S, Kumar A, Trivedi SS, Puri M, Sarin SK. Maternal and Fetal outcome in pregnant women with acute hepatitis E virus infection. Ann Intern Med. 2007;147(1):28–33.

50. ProMED-mail 11th Aug 2017.https://www.thecable.ng/hepatitis-e-kills-four-pregnant-women-borno-400-idps-infected

51. Temitope Oluwasegun Cephas Faleye, Moses Olubusuyi Adewumi, Ijeoma Maryjoy Ifeorah, Ewean Chukwuma Omoruyi, Solomon Adeleye Bakarey, Adegboyega Akere, Funmilola Awokunle, Hannah Opeyemi Ajibola, Deborah Oluwaseyi Makanjuola, and Johnson Adekunle Adeniji. Detection of hepatitis B virus isolates with mutations associated with immune escape mutants among pregnant women in Ibadan, southwestern Nigeria.Springerplus. 2015; 4: 4

52. WHO Position Paper. Hepatitis E Vaccine. weekly Epidemiological records.1st May 2015;90(18):185–200. http://www.who.int/wer/2015/wer9018/en/

